# Deacetylation of IFIT2 mediated by HDAC5 promotes the stemness and progression of glioma

**DOI:** 10.1101/2021.04.04.438236

**Authors:** Ying Liu, Kun Zhang, Xingzhi Peng, Zhuan Zhou, Peijun Zhou, Siyuan Tang, Dan Li, Liangfang Shen, Deyun Feng, Lifang Yang

## Abstract

**Background:** Glioma is the most common primary brain tumor, and the tumor stemness is a major regulatory factor affecting the progression, metastasis and recurrence of glioma. Recent research has shown that, nonhistone acetylation is widely involved in key cellular processes, including stemness regulation. The deacetylase inhibitors are promising new drugs, but their application and molecular mechanism in glioma have not been elucidated.

**Methods:** CCK8 and colony formation assay were used to detect cell proliferation, transwell assay was used to detect cell migration, flow cytometry was used to analyze cell apoptosis, sphere formation assay and western blot were used to detect the status of stemness. RNA-sequence, quantitative PCR and western blot were performed to screen the key molecules mediating LBH589 function. Immunoprecipitation (IP) and western blot were used to analyze the acetylation level of IFIT2. The SiRNA target HDAC4 or HDAC5, overexpression plasmids of acetyltransferases were used to identify the acetyltransferase and deacetylase regulating IFIT2. The regulatory mechanism was explored by IP and ubiquitination analysis. Finally, the xenograft tumor model in nude mice was constructed and further analyzed in vivo.

**Results:** The data showed that IFIT2 mediates the HDACi LBH589 inhibition on cell proliferation, migration and stemness, and contribution to autophagy and apoptosis in glioma. And the down-regulation of IFIT2 in glioma was confirmed to be related to its deacetylation by overexpression HDAC5, which promotes the stemness and progression of glioma. Further, deacetylation of IFIT2 by HDAC5 was demonstrated to induce its ubiquitination and subsequent protein instability, which led to loss of anti-tumor activity for IFIT2, and acceleration to glioma stemness and progression. In addition, the results indicated that IFIT2 inhibits PKC pathway, and suppressing of IFIT2 promotes tumor growth in vivo.

**Conclusions:** These results not only clarify a novel post-transcriptional regulatory mode of IFIT2, but also provide a new sight of molecular mechanism for HDACi in glioma.

## Introduction

Glioma, one of the common primary brain tumors, takes accounts for 80% of adult primary malignancy in central nerve system [1]. Annual new cases of glioma is 38 to 48 per million worldwide, with a high morbidity rate of 7% and mortality rate of 5.5% of all cancers, posing a serious threat to all human beings [2]. Despite of combined therapies including surgery, radiation and chemotherapy in glioma, the median survival of glioma is still less than 2 years as therapy resistance frequently occurs. The therapy resistance of glioma was mainly due to the inherent characteristics of cells and tumor microenvironment, which may be attributed to the existence of stemness [3]. Tumor stemness refers to characteristics of self-proliferation and pluripotency in cancer cell, which always results in tumor initiation, progression, metastasis and relapse [4]. Therefore, exploration of tumor stemness and the underlying regulation mechanism may provide us with new interferences and promising targets in glioma clinical utility.

Epigenetics, defined as heritable changes in gene expression but without alterations in the nucleotide sequence, consist of DNA methylation, histone modifications, chromatin remodeling and non-coding RNAs, playing a fundamental role in many biological processes including tumorigenesis and tumor progression [5]. Histone acetylation is a vital modification in epigenetics, which is dynamically mediated by histone acetyltransferase (HATs) and deacetylase (HDAC). In general, histone acetylation inhibits the coalition of DNA and histone, opens the chromatin structure and makes it accessible to transcription factors, thus activating gene transcription [6]. For instance, tumor suppressor gene ARID1A recruits HDAC1 to the promoter of stemness marker ALDH1A1 and reduces histone H3K27 acetylation, silencing the expression of ALDH1A1 and thereby inhibiting the metastasis of bile duct cancer cells [7]. HDAC11 decreased H3K9 acetylation in promoter of LKB1, enhancing the stemness and progression of liver cancer through LKB1 / AMPK pathway [8]. Hence, histone acetylation is closely connected to tumor progression on the basis of cancer stemness.

Considering that most tumor suppressor genes were silenced in cancer because of the modified acetylation status, histone deacetylase inhibitors (HDACi) were developed as a new class of anti-tumor drugs in recent years and reported to function in many tumors including glioma [9, 10]. Interestingly, acetylation was also found to occur in nonhistone proteins, involving in regulations of enzyme activity, protein stability and protein-protein interactions. Nonhistone acetylation is associated with various cellular processes such as transcription, signal transduction, DNA damage repair, autophagy and metabolism, which consequently contributed to tumor growth, metastasis, stemness, drug resistance and relapse [11]. High acetylation of E3 ubiquitin ligase MDM2, for example, ensured recruitment of de-ubiquitination enzyme HAUSP to P300 and inhibited self-degradation of P300, thereby more MDM2 translocated from the nucleus to the cytoplasm, which finally led to the increasing ubiquination of P53 and regression of P53-dependent apoptosis [12].Besides, acetylation of K403 in glucose-6-phosphate dehydrogenase (G6PD) restrained its dimerization and further inactivated G6PD, but SIRT2 reversed such progress through deacetylation of G6PD and consequently stimulated the production of NADPH in pentose phosphate pathway to resist oxidative damage [13]. All of these studies indicate that direct acetylation of nonhistone is essential to tumor progression.

Interferon inducible protein 2 (IFIT2), also known as ISG54, is related to the function of cytoskeleton. Intensive studies have shown that IFIT2 downregulated TNF-α, Akt, and PKC signaling pathways to decline cell proliferation, migration and inhibit cell apoptosis in many cancers like oral squamous cell carcinoma (OSCC) and chronic myeloid leukemia (CML) [14-16]. A recent research domonstrated that IFIT2 may involve in tumor stemness since baicalein was found to sensitize radio-chemotherapy and inhibit the stem-like properties including cell sphere formation and stemness marker OCT3/4 and ABCG2 expression in human breast cancer cells [17].

In present study, we investigated the role of IFIT2 mediating the HDACi LBH589 inhibition on cell proliferation, migration and stemness, and contributions to autophagy and apoptosis in glioma. We further revealed that deacetylation of IFIT2 by HDAC5 induced its ubiquitination and subsequent protein instability, which led to loss of anti-tumor activity and acceleration to glioma stemness and progression.

## Materials and Methods

### Cell cultures and reagents

Normal glial cells HEB and glioma cells HS683(HTB-138), SHG44,SW1088 (HTB-12),U251MG, SF-295, SF-126 and HEK293T(ATCC®CRL-11268™) were maintained in DMEM (Gibco BRL, Grand Island, NY, USA) supplemented with 10% fetal bovine serum(Gibco BRL) at 37 °C in 5% CO_2_. Histone deacetylase inhibitors LBH589 (S1030), LMK-235(S7569), MS-275(S1053), TSA(S1045) and NAM(S189), cycloheximide (CHX, S7418) were purchased from Selleck Chemicals (Houston, TX,USA), DMSO (D2650) were purchased from Sigma-Aldrich (St. Louis, MO, USA), MG-132(HY-13259) were purchased from MCE (Shanghai, CHN).

### Plasmid, siRNA and transfection

pcDNA3.1-3xFlag-IFIT2 was purchased from Addgene (#53555). The constructs His-Ubiquitination was a gift from Prof. Gang Huang (Cincinnati Children’s Hospital, Cincinnati, OH, USA). MYC-P300 and CBP is a gift from Prof. Jiuhong Kang (School of Life Science and Technology, Tongji University, Shanghai, CHN). HA-TIP60, HA-PCAF and HA-GCN5 purchased from Youbio company (Changsha, CHN). All SiRNA were synthesized in the Ribobio company (Guangzhou, CHN). Sequences: Si-IFIT2#1 5′-CGACTCTCAGACGTTCAGATT -3′; Si-IFIT2#2 5′-GCATTCCTTCAGGAGCTGAAT -3′; Si-IFIT2#3 5′-GCAACCTACTGGCCTATCTAA -3′; Si-HDAC4#1 5′-GGGAATGTACGACGCCAAA -3; Si-HDAC4#2 5′-TGTCGACCTCCTATAACCA-3′; Si-HDAC4#3 5′-AGCCCATTGAGAGCGATGA -3′; and Si-HDAC5#15′-GGACTGTTATTA GCACCTT -3′; Si-HDAC5#2 5′-CAACGGGAACTTCTTTCCA -3′; Si-HDAC5#3 5′-CAACGGGAACTTCTTTCCA -3′. According to the manufacturer’s instructions, indicated plasmids and SiRNAs were transfected using Lipofectamine 3000 (L3000015, Invitrogen Carlsbad, CA, USA).

### Cell viability assay

Cells were seeded in 96-well plates (4.0 × 10^3^ cells per well). Cell viability was measured by adding 10% CCK8 (C0005,Targetmol, Shanghai, CHN) at different times (0, 24, 48, 72 hrs) for an additional 1 h. Absorbance was measured at 450 nm.

### Colony formation assay

Cells were seeded in 6-well culture plates and treated with 0.075 μM LBH589. Cells were then harvested and counted under microscope. 500 cells/well were added in 2 mL medium in 6-well culture plates and cultured for 10 days. Next, cells were fixed by 4% paraformaldehyde (BL539A, BioSharp Hefei, China) for 15 min and stained by crystal violet for 15 min. After staining, cells were photographed under a conventional microscope (DMI3000B, Leica) and counted manually.

### Transwell invasion assay

The cell invasion assays were conducted using PET-track-etched membrane invasion chamber (BD Biosciences, San Jose, CA, USA) with 20 μL of diluted Matrigel (356234, BD Biosciences). 4 × 10^4^ cells were added to the upper chamber without FBS. The lower chamber was filled with 1 ml of DMEM with 20% FBS. The chambers were incubated in 37 °C for 24 h, and the invaded cells were stained with crystal violet. The cells that adhered to the membranes were imaged under a conventional microscope (DMI3000B, Leica) and counted manually.

### Flow cytometry assay

Cells were seeded in 6-well culture plates, then trypsinized and washed with cold PBS for 3 times. Annexin V-FITC/PI apoptosis kit (KGA108-1, KeyGene Biotech, Jiangsu, CHN) were used in this study according to the manufacturers’ instruction. The apoptotic analysis was performed with a flow cytometer (FlowSight, Millipore, Billerica, MA, USA).

### Sphere formation assay

Cells were seeded into ultralow-attachment 6-well plates (Corning, NY, USA) with serum-free DMEM/F12 medium containing EGF (PeproTech, NJ, USA), basic Fibroblast Growth Factor (bFGF, Sigma-Aldrich), B27 Supplement and N-2 Plus Media Supplement (Life Technologies, NY, USA). After incubation for 1-2 weeks, cells were counted and photographed under a conventional microscope (DMI3000B, Leica).

### Real-time quantitative PCR (QPCR)

Total RNA was extracted from cells and quantified using RNA Isolation Reagent kit (9109, Takara, Tokyo, Japan). cDNA was obtained from 1 µg of total RNA from each sample (Super M-MLV reverse transcriptase, TAKARA) and QPCR was performed using TaqMan™ Gene Expression Master Mix (4369016, Thermo Fisher Scientific Waltham, MA, USA) on the ABI 7500 QPCR System (Foster City, CA, USA). For each sample and experiment, triplicates were made and normalized by β-actin mRNA levels. The primers: IFIT2 forward, 5′-AAGGAAAAAA GCGGCCGCATGCGCCATGAGTGAGAACAATA -3′, reverse, 5′-CGGGATCCTTATTTCCCCATTCCAGCTTGAT -3′; P300 forward,5′-GTTCCTTCCTCAGACTCAGTTC -3′, reverse, 5′-CATTATAGGAGAGTTCACCGGG -3′; β-actin forward, 5′-CATGTACGTTGCTATCCAGGC-3′, reverse, 5′-CTCCTTAATGTCACGCACGAT-3′.

### RNA sequencing

SW1088 and U251MG cell with DMSO and LBH589 treatment were RNA sequenced by Novogene company (Beijing, China) for transcriptome sequencing and analysis.

### Western blot and Immunoprecipitation (IP) assays

For western blot analysis, cells were lysed with IP lysis buffer supplemented with a protease inhibitor ‘cocktail’. Protein concentrations in the extracts were measured by BCA assay (AR1189A, BOSTER, Wuhan, CHN). Equal amounts of extracts were separated by SDS-PAGE, then transferred onto polyvinylidene fluoride membrane (IPVH00010, Merck Millipore, Billerica, MA, USA), blocked with 5% dry nonfat milk in Tris-buffered saline (pH 7.4) containing 0.1% Tween-20 and probed with the antibody for western blot analysis and the membrane was stained with the corresponding primary and secondary antibodies. The specific bands were analyzed with an Moleculai Imager® Chemidoc™ XRS+ with Image Lab™ Software imaging system (BIO-RAD, Hercules, CA, USA).

For immunoprecipitation (IP), whole-cell extracts were lysed in IP Lysis Buffer (20118ES60, Yeasen, Shanghai, CHN) and a protease inhibitor ‘cocktail’ (B14001, Bimake, Shanghai, CHN). Cell lysates were centrifuged for 15 min at 13,000 × g. Supernatants were collected and incubated with protein G magnetic beads (10004D, Invitrogen) together with specific antibodies. After overnight incubation, protein G magnetic beads were washed five times with IP wash buffer. IP was eluted by SDS-PAGE loading buffer (CW0027S, CWBiO, Beijing, CHN). The following primary antibodies were used: Anti-Nestin (33475S, 1:1000), Anti-Nanog (4903T, 1:1000), Anti-LC3 (3868S,1:1000), Anti-Acetylated-Lysine Antibody (9441,1:1000 for western blot, 1:200 for IP), Anti-Acetyl-Histone H3 (Lys27, 8173, 1:1000), Anti-P-PKC (2060S, 1:1000), anti-ubiquitin (3936), and Anti-P300 (86377, 1:1000) were purchased from CST (Danvers, MA,USA), Anti-CD133(ab216323, 1:1000) were purchased from Abcam (Cambridge, MA, USA), Anti-Oct4 (AF2506, 1:500) were purchased from Beyotime Biotechnology (Jiangsu, CHN), Anti-P62 (SC-25575, 1:1000) were purchased from Santa Cruz Biotechnology (Santa Cruz, CA, USA), Anti-actin (ab0261,1:10000) were purchased from Abconal (Wuhan, CHN), Anti-IFIT2(612604-1-AP, 1:300 for western blot, 1:100 for IP), Anti-HDAC4(617449-1-AP, 1:500), Anti-HDAC5 (616166-1-AP, 1:200), Anti-MYC (616286-1-AP, 1:3000), Anti-AKT(660203-2-IG, 1:1000), Anti-P-AKT(66444-1-IG, 1:1000) were purchased from Proteintech (Wuhan, CHN), Anti-Flag (F1804, 1:1000 for western blot, 1:500 for IP) were purchased from Sigma-Aldrich (St. Louis, MO, USA).

### Bioinformatics analysis

The gene expression profiles and survival curves of the relationship between the expression of IFIT2, HDAC5 and P300 were from the Chinese Glioma Genome Atlas (CGGA) and Cancer Genome Atlas Program(TCGA).

### Animal Experiments

Animal experiments were conducted with the approval of the Institutional Animal Care and Use Committee of the Xiangya School of Medicine of Central South University and conform to the legal mandates and federal guidelines for the care and maintenance of laboratory animals. Ten 5-week-old male athymic nude C57/BL6J mice were housed under pathogen-free conditions. U251MG cells (1 × 10^7^ cells in 150 μL PBS) were injected subcutaneously into the right flank. The tumor volume was measured using a vernier caliper every 3 days and tumor volume was calculated as below: V (cm^3^) = width^2^ × length /2 [18]. When the tumor volume reached 60–80 mm^3^, a half of mice were injected intraperitoneally with LBH589 (20 mg/kg/day), the others were injected intraperitoneally with DMSO. After injection for 4 weeks, animals were killed and tumor volume and weight were recorded. The tumor tissues were fixed with 10% buffered formalin and embedded in paraffin for the immunohistochemical experiments.

### Immunohistochemistry (IHC)

Tissue sections were deparaffinized in xylene and rehydrated with ethanol, and then preincubated with 10% normal goat serum in PBS (pH 7.5) followed with incubation with primary antibody overnight at 4°C, and then stained with biotinylated secondary antibody (SAB4600042, Sigma-Aldrich) for 1 h at room temperature, incubated with reaction enhancement solution for 20 minutes. The peroxidase reaction was developed with DAB kit (ZLI-9019, ORIGENE, Beijing, CHN) and the slides were counterstained with hematoxylin. The concentration of primary antibody was IFIT2 (612604-1-AP, Proteintech,1:100) and HDAC5 (616166-1-AP, Proteintech,1:100). The tissues were photographed under a conventional microscope (DMI3000B, Leica).

### Statistical analysis

All the statistical data are presented as mean ± S.E.M., statistical significance of the differences was determined using the Student *t* test. *p<* 0.05 was considered statistically significant.

## Results

### LBH589 induces apoptosis and autophagy and inhibits the proliferation, metastasis and stemness of glioma cells

To determine the effect of histone deacetylase inhibitor LBH589 on glioma cells, firstly, CCK8 assay was used to analyse the proliferation ability of glioma cell lines, which treated with serial dilutions of LBH589 (0.025, 0.05,0.075 μM) at different times (24, 48, and 72 hrs), the results showed that LBH589 significantly inhibited the proliferation of glioma cell SW1088 and U251MG in a concentration dependent manner (p<0.001) (Figure 1A). Also, similar results were obtained in the colony formation assay, indicating that LBH589 notably repressed the proliferation in glioma cells (Figure 1B). Further, transwell migration assays showed that LBH589 treatment significantly decreased migration ability in both SW1088 and U251MG cells compared with DMSO treatment (p<0.01) (Figure 1C). Furthermore, flow cytometric analysis showed that LBH589 treatment induced apoptosis of SW1088 and U251MG (p<0.01) (Figure 1D). Moreover, the western blot data exhibited that LBH589 could elevated the expression of LC3 and deceased the level of P62, which indicated that LBH589 could induced autophagy in glioma (Figure 1E). In addition, LBH589 could impair the sphere formation of glioma cell (p<0.05) (Figure 1F). And stem cell markers such as Nestin, CD133, Nanog, Oct4 were consistently downregulated in both SW1088 and U251MG which treated with LBH589 (Figure 1G). Collectively, these results indicate that LBH589, the histone deacetylase inhibitor, could inhibit the proliferation, metastasis and stemness of glioma cells, and induce apoptosis and autophagy.

**Fig.1.**
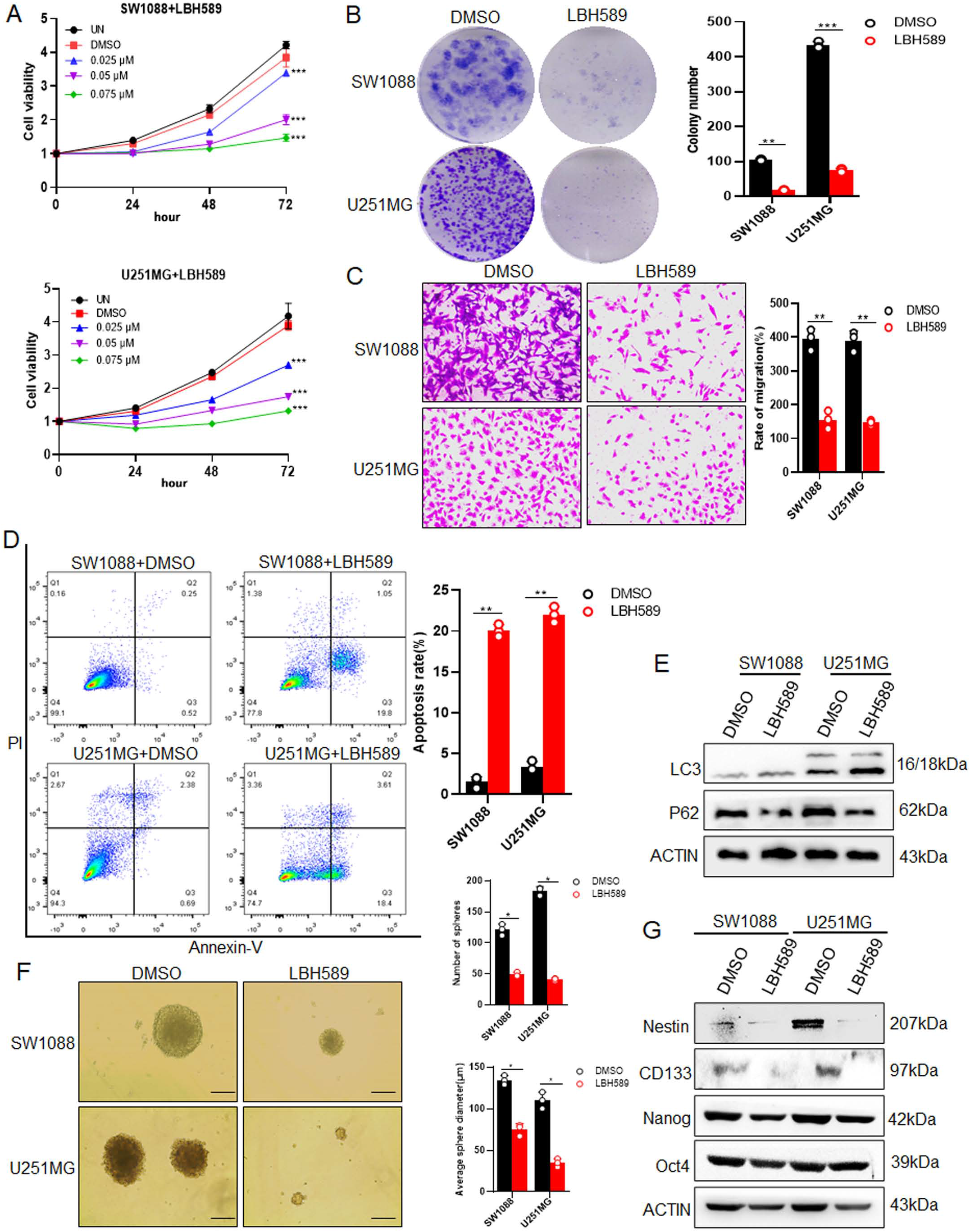
LBH589 induces apoptosis and autophagy and inhibits the proliferation, metastasis and stemness of glioma cells. (**A)** SW088 and U251MG cells were treated with 0.025, 0.05, and 0.075 μM of LBH589 for 24, 48 and 72 hrs, cell viability was determined using the CCK8 assay. **(B)** The colony numbers in soft agar (2 weeks) and **(C)** the invasion numbers of cells after treatment with 0.075 μM LBH589 are shown with histograms. **(D)** Flow cytometric analysis were performed to observe cell apoptosis after treatment with LBH589 for 48h. **(E)** Western blot were analyzed to detect LC3 and P62 in SW1088 and U251MG cells after treatment with LBH589. **(F)** The tumor sphere formation assay was performed in FBS-free medium in low-density cultures per 200 cells with six replicates, the number and size of tumor spheres formed by SW1088 and U251MG cells was calculated after treatment with LBH589. **(G)** The differential expression of several stemness-related genes, including Nestin, CD133, Nanog and Oct4 in SW1088 and U251MG cells was validated by western blot after treatment with LBH589. Scale bar: 50 μm, UN: untreated, Columns: mean of three replicates; *p<0.05, **p<0.01, and ***p<0.001

### IFIT2 is responsible for LBH589 mediated biological function in glioma

To identify the molecule of LBH589-mediated biological function in glioma cells, the transcriptome changes of SW1088 and U251MG cell were analyzed after treatment with LBH589 by RNA-Seq. Then the transcription levels of multiple tumor suppressor genes were selected for further study. The data suggested that IFIT2 was the most obviously upregulation gene in SW1088 and U251MG cells treated with LBH589 (Figure 2A). IFIT2 were detected by QPCR and western blot in glioma cell, and the results revealed its significant down-regulation in glioma cells, compared with normal glial cells HEB (p < 0.001) (Figure 2B,2C). Similarly, The CGGA and TCGA analysis showed lower IFIT2 in glioma compared with non-tumor tissues (p < 0.05) (Figure S1A), and the expression of IFIT2 in high-grade glioma was lower than that in low-grade glioma (p < 0.001) (Figure S1B). Interestingly, the results of both mRNA (Figure 2D) and protein levels (Figure 2E) was observed that IFIT2 was up-regulated in a time dependent manner. Further, overexpression of IFIT2 in SW1088 and U251MG cells significantly inhibited cell proliferation (p < 0.001) (Figure S2, Figure 2F) and induced apoptosis (p < 0.001) (Fig. 2G), meanwhile, reduced the expression of Nestin, CD133, Nanog and Oct4 (Figure 2H).

**Fig.2.**
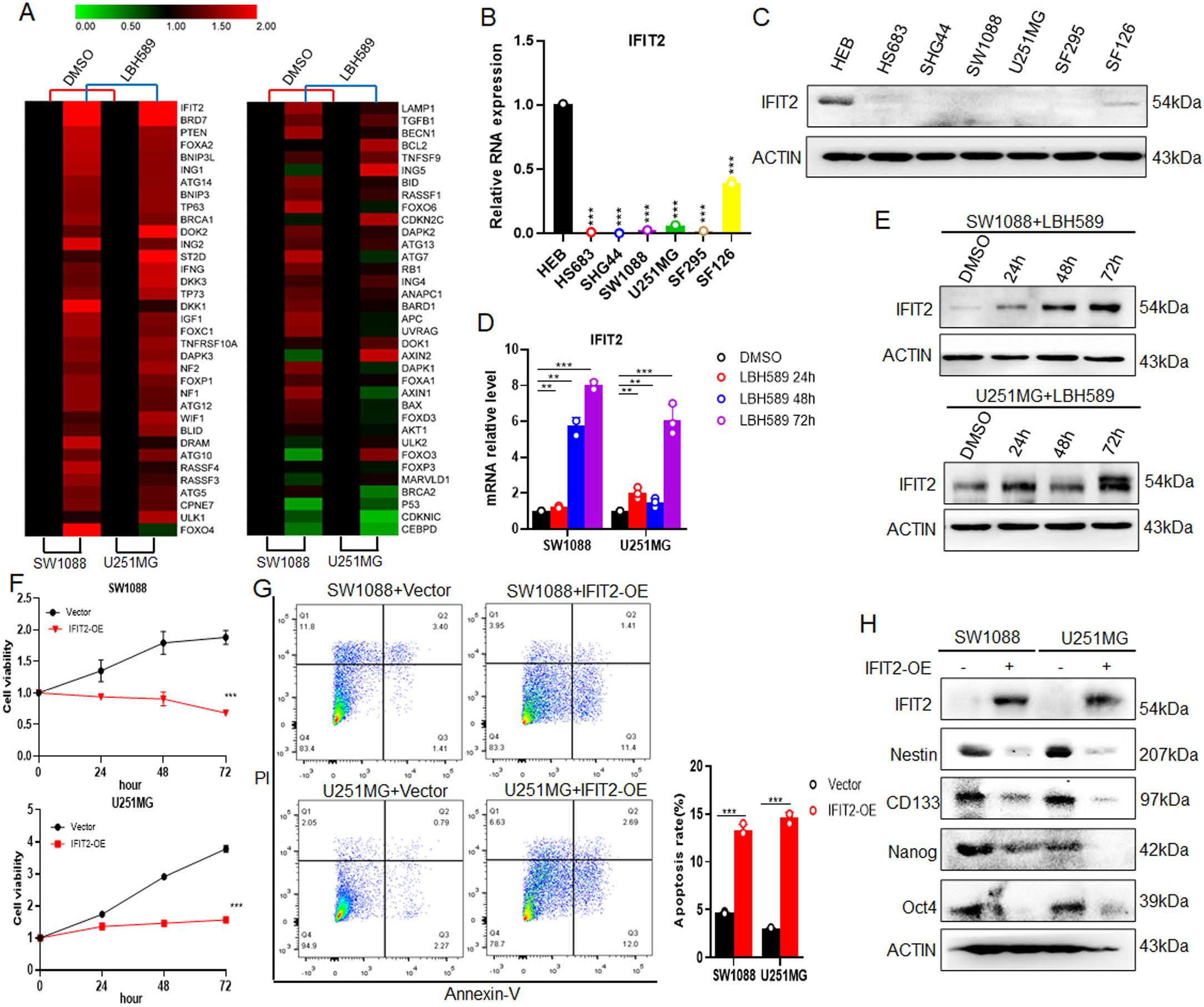
LBH589 upregulates IFIT2 expression. **(A)** RNA-Seq was performed to analyze alteration of tumor suppressor genes in SW1088 and U251MG cells after treatment with LBH589. **(B)** QPCR and **(C)** Western blot were performed to detect IFIT2 expression in HEB and glioma cells. **(D)** QPCR and **(E)** Western blot were performed to detect IFIT2 expression in glioma cells with LBH589 treatment for different times. **(F)** the CCK8 assay and **(G)** Flow cytometric analysis was performed to observe glioma cell apoptosis. **(H)** The expression of IFIT2, Nestin, CD133, Nanog and Oct4 was analyzed by western blot after transfected with overexpression IFIT2 plasmid. Columns: mean of three replicates; **p<0.01, and ***p<0.001

Furthermore, combination of LBH589 and IFIT2 siRNA synergistically showed that knockdown of IFIT2 expression significantly elevated the proliferation of glioma cells stimulated by LBH589 (p < 0.001) (Figure S3, Figure 3A). Thus, the promotion of apoptosis by LBH589 was strongly suppressed by the low expression of IFIT2 (Figure 3B). Moreover, IFIT2 inhibition significantly increased the transwell migration of SW1088 and U251MG cells after treatment of LBH589 (p < 0.05) (Figure 3C). Similar effect was observed by tumor sphere formation assay that disruption of IFIT2 by siRNAs strongly rescued the suppression of tumor sphere formation by LBH589, as revealed by the decreased number and size of tumor spheres (Figure 3D). Notably, the knockdown of IFIT2 significantly attenuated the level of autophagy and inhibition of LBH589 on stemness makers by LBH589 in glioma cells (Figure 3E). These results reveal that IFIT2 is responsible for LBH589 mediated biological function in glioma.

**Fig.3.**
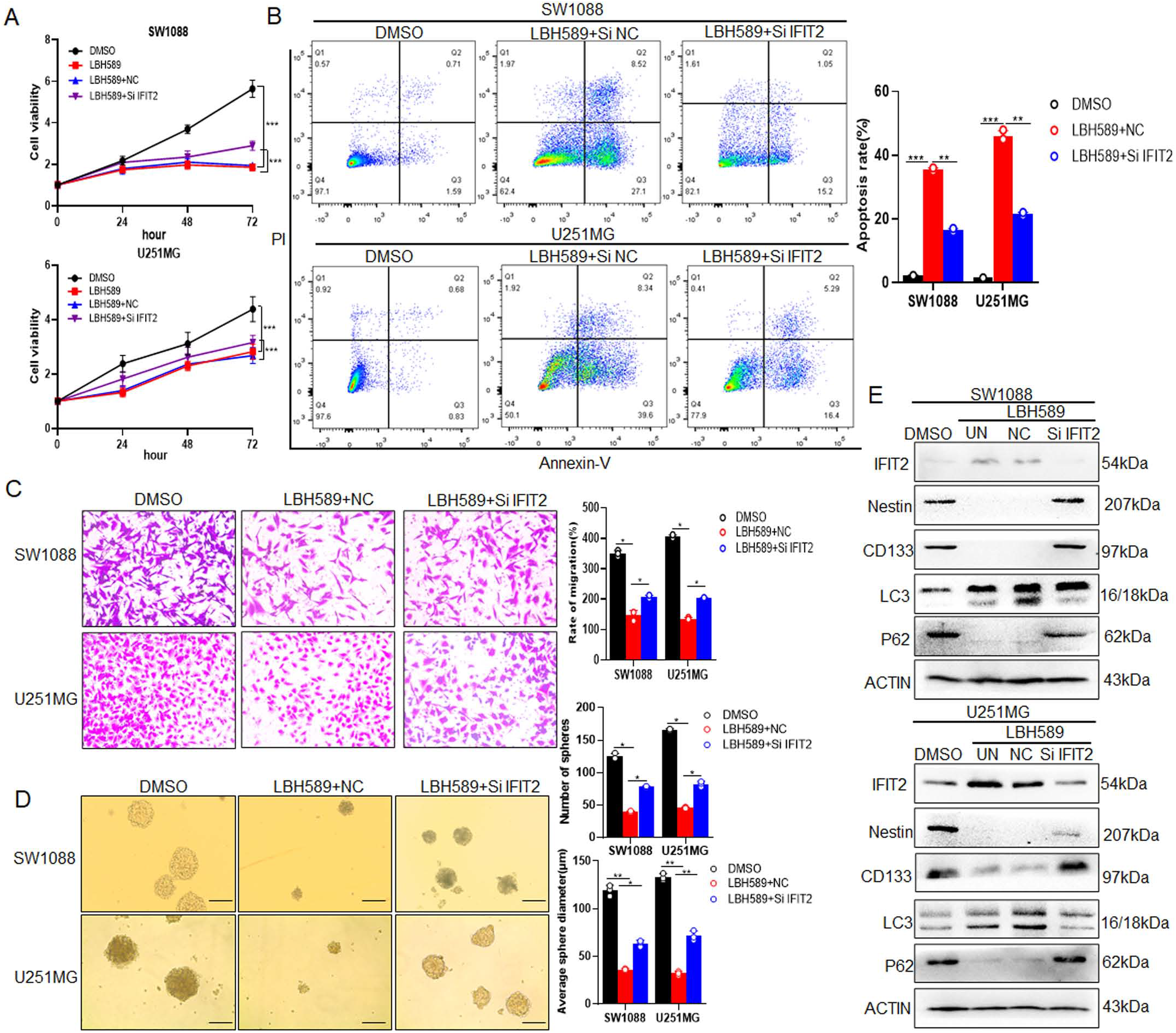
IFIT2 is responsible for LBH589 mediated biological function in glioma. After transfection with SiRNA-NC or SiRNA-IFIT2 for 48h, **(A)** the viability of cells after treatment with LBH589 was assessed by the CCK8 assay. **(B)** Flow cytometric analysis were performed to observe cell apoptosis after treatment with LBH589 for 48h. **(C)** Matrigel invasion assay in cells treated with LBH589. The number of cells per field invading onto the lower surfaces of the filter was counted. **(D)** The number and the size of tumor spheres formed by SW1088 and U251MG cells after treatment with DMSO or LBH589 was calculated. **(E)** The expression of Nestin, CD133, LC3, P62 and IFIT2 in SW1088 and U251MG cells after treatment with LBH589 was detected by western blot. UN: untreated, NC: negative control. Columns: mean of three replicates; *p<0.05, **p<0.01, and ***p<0.001

### IFIT2 is deacetylated by HDAC5

To clarify the up-regulation mechanism of IFIT2, the histone acetylation after LBH589 was detected. The data showed that histone H3 acetylation at lysine 27 (H3K27Ac) was most significantly up-regulated (Figure S4), suggesting that LBH589 increased IFIT2 mRNA expression through H3 acetylation, which is consistent with the previous studies [19]. Meanwhile, we analyzed the acetylation of endogenous IFIT2 in U251MG and SW1088 cells (Figure 4A) and in 293T cell with IFIT2 exogenous expression (Figure 4B). Using a specific antibody against acetylated lysine, the data indicated the strong acetylations of IFIT2, suggesting that IFIT2 was acetylated directly. Next, we identified the potential deacetylases of IFIT2 in SW1088 and U251MG cells. Treatment with trichostatin A (TSA), a broad spectrum inhibitor of HDAC (class I and II), instead of NAM, an inhibitor of SIRT (class III), increased IFIT2 expression in the cells (Figure S5). Then, treated cells with LMK-235, a specific inhibitor HDAC4/HDAC5 [20], and MS-275, the effect of the class Ⅰ HDAC [21], the result showed that LMK-235 could up-regulate the IFIT2 expression (Figure 4C). Further, western blot were used to detect the expression of IFIT2 by transfecting with SiRNA against HDAC5 rather than HDAC4 (Figure 4D). Furthermore, IP analysis identified a specific association between IFIT2 and deacetylase HDAC5 in glioma SW1088 cells, and the association was weakened after knocking down HDAC5 (Figure 4E). Moreover, the interaction of HDAC5 and IFIT2 greatly increased the IFIT2 acetylation, and knockdown of HDAC5 also raised the acetylation level of IFIT2 in glioma cells (Figure 4F, 4G). Taken together, these results suggestted that HDAC5 interacts directly with IFIT2, which mediates the deacetylation of IFIT2.

**Fig.4.**
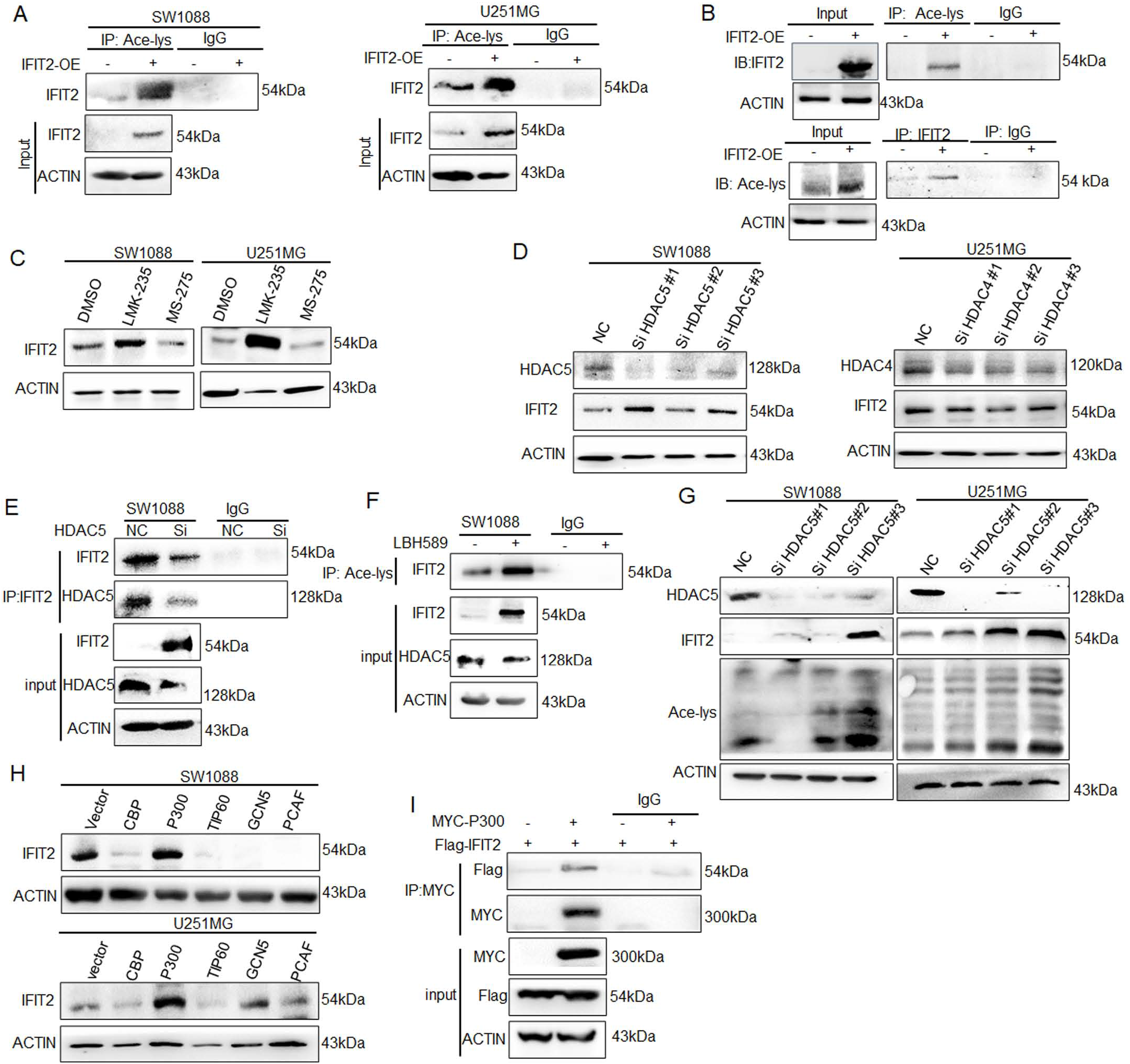
IFIT2 is deacetylated by HDAC5. **(A)** Acetylation of Flag-IFIT2 in SW1088 and U251MG cells was Immunoprecipitated with anti-acetyl-lys, and the precipitates were analyzed using an anti-IFIT2 antibody (Ace-lys). **(B)** Acetylation of IFIT2 in 293T cells was analyzed by IP with an anti-acetyl-lys or anti-IFIT2 antibody, then followed by western blotting for anti-IFIT2 or anti- Ace-lys. (C) SW1088 and U251MG cells were treated with LMK-235 and MS-275 for 48 hrs for western blotting analysis. (D) After transfection with siRNA-NC or SiRNA-HDAC4 or SiRNA-HDAC5, the expression of IFIT2 in SW1088 and U251MG cells was detected by western blot. **(E)** The interaction between IFIT2 and HDAC5 in SW1088 cell was measured by IP using IFIT2 and HDAC5 antibody, respectively. **(F)** After treatment with DMSO or LBH589, IP in SW1088 and U251MG cells with anti-IFIT2 were confirmed by western blot with acetyl-lys antibody. **(G)** After transfection with SiRNA-NC or SiRNA-HDAC5, the expression of IFIT2, anti-acetyl-lys and HDAC5 in SW1088 and U251MG cells was detected by western blot. **(H)** After transfection with different acetyltransferases, the expression of IFIT2 in SW1088 and U251MG cells was detected by western blot. **(I)** After transfection with MYC-P300 or Flag-IFIT2, IP in 293T cells with anti-MYC were confirmed by western blot with anti-Flag antibody.

Histone acetylation is regulated by histone acetyltransferases (HATs) and histone deacetylases (HDACs).We found that only P300 could promote the acetylation of IFIT2. In contrast, other acetyltransferases, including CBP, TIP60, GCN5 and PCAF, failed to acetylate IFIT2 in SW1088 and U251MG cells (Figure 4H). After transfecting with MYC-P300 and FLAG-IFIT2 in 293T cells, and the data indicated a combination between P300 and IFIT2 (Figure 4I). These results indicated that HDAC5/P300 is involved in the acetylation of IFIT2.

### Acetylation promotes the stability of IFIT2 via inhibiting its ubiquitination

The previous studies reported that acetylation modification could inhibit protein stability, and the mechanism of acetylation-dependent protein stability was prevention of protein ubiquitylation, thereby inhibition of proteasome-dependent degradation [22, 23]. Inhibition of protein synthesis with cycloheximide (CHX) showed that knockdown of HDAC5 could significantly extend the half-life of IFIT2 protein (Figure 5A). Further, treating the cells with MG-132 to inhibit proteasome activity, the results showed that MG-132 exposure enhanced the up-regulation IFIT2 induced by HDAC5 silence (Figure 5B). Meanwhile, the data also indicated that LBH589 promoted the IFIT2 stability and increased the IFIT2 expression after treatment with CHX and MG-132 (Figure 5C, 5D), supporting that HDAC5 enhances the protein stability and stimulation of IFIT2. To clarify whether the regulation of HDAC5-mediated IFIT2 acetylation depending on its ubiquitination, we co-transfected Si-HDAC5, IFIT2 overexpression plasmid and Ub plasmid in 293T cells, and the co-IP data showed that depletion of HDAC5 significantly decreased the ubiquitination of IFIT2 (Figure 5E). These data indicate that HDAC5-mediated deacetylation could down-regulate the protein stability and expression of IFIT2 through ubiquitination modification.

**Fig.5.**
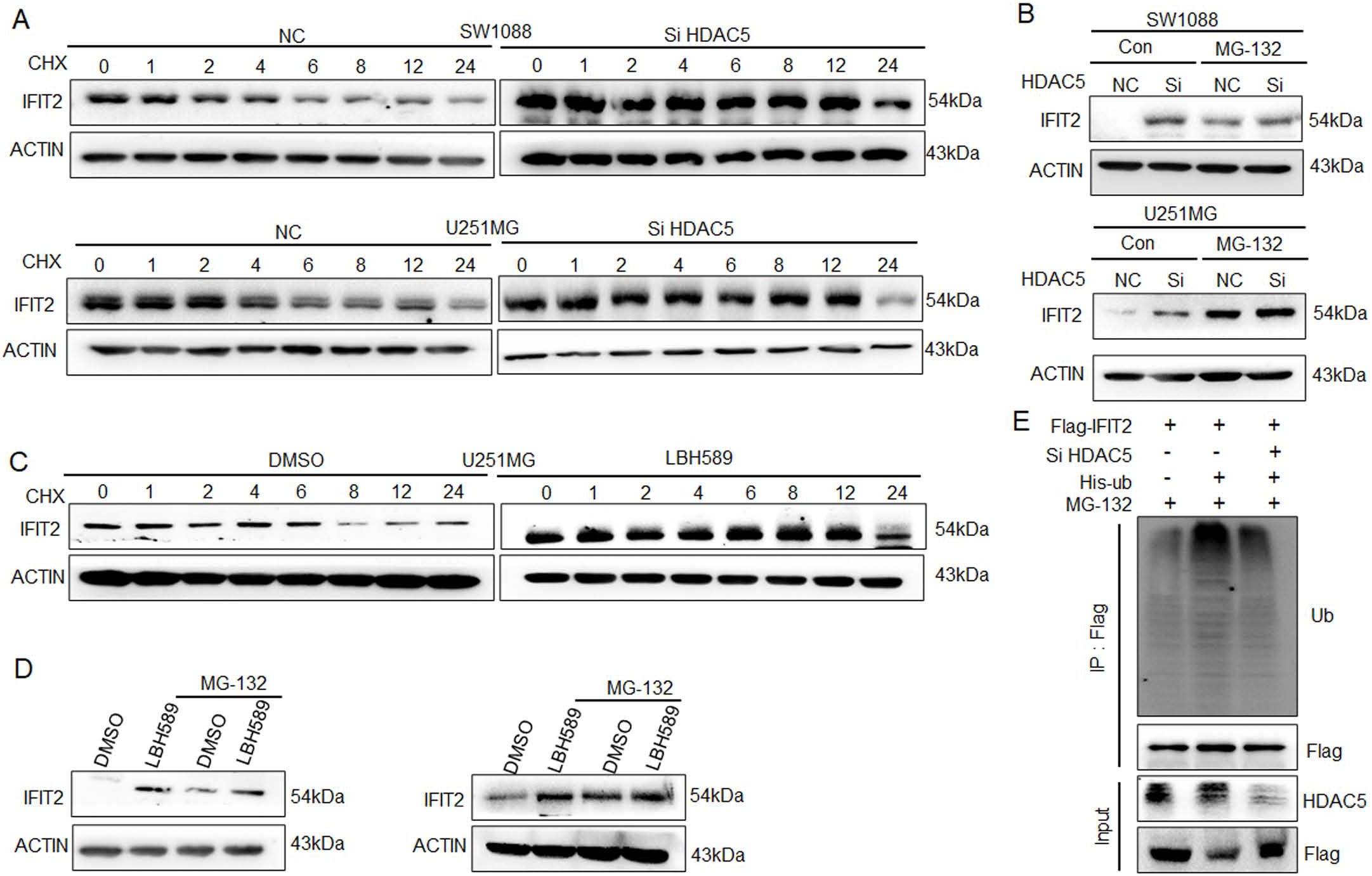
Acetylation promotes the stability of IFIT2 via inhibiting its ubiquitination. SW1088 and U251MG cells with SiRNA-NC or SiRNA-HDAC5 transfection for 48h were treated with CHX **(A)** and MG-132 **(B)**, the protein expression of IFIT2 was measured. SW1088 and U251MG cells with LBH589 treatment were exposed to CHX **(C)** and MG-132 **(D)**, the protein expression of IFIT2 was measured. **(E)** After transfection with SiRNA-NC or SiRNA-HDAC5, His-Ubiquintination and Flag-IFIT2, the ubiquintination in 293T cells was analyzed by western blot using anti-Ubiquitin antibody.

### Overexpression of HDAC5 promotes the stemness and progression of glioma

To explore the biological role of HDAC5 in glioma, firstly, the HDAC5 expression was detected, and the results revealed that HDAC5 was commonly overexpressed in the glioma cells. Similarly, the CGGA data showed that HDAC5 was highly expressed in glioma compared to normal brain tissue, and its expression was positively correlated with tumor grade (Figure S6A, S6B). Besides, the patients with high express of HDAC5 had a significantly poor prognosis (p<0.001) (Figure S6C). The P300 expression was examined using QPCR and western blot assay as well, and the data showed that P300 was low expression in glioma cells compared with HEB cells (p<0.001) (Figure S7A, S7B), and the patients with low express of P300 had a significantly poor prognosis (p<0.001) (Figure S7C, S7D). To investigate the role of HDAC5 in glioma, HDAC5 was knocked down by SiRNA in SW1088 and U251MG cells. The data showed that knockdown of HDAC5 inhibited cell growth (p<0.001) (Figure 6A), induced cell apoptosis (p<0.01) (Figure 6B) and decreased the cell migration (Figure 6C). Sphere formation assay results indicated that knockdown of HDAC5 inhibited the stemness in SW1088 and U251MG cells (Figure 6D), and decreased the expression of Nestin, CD133 and Oct4. These results demonstrated that overexpression of HDAC5 promotes the stemness and progression of glioma.

**Fig.6.**
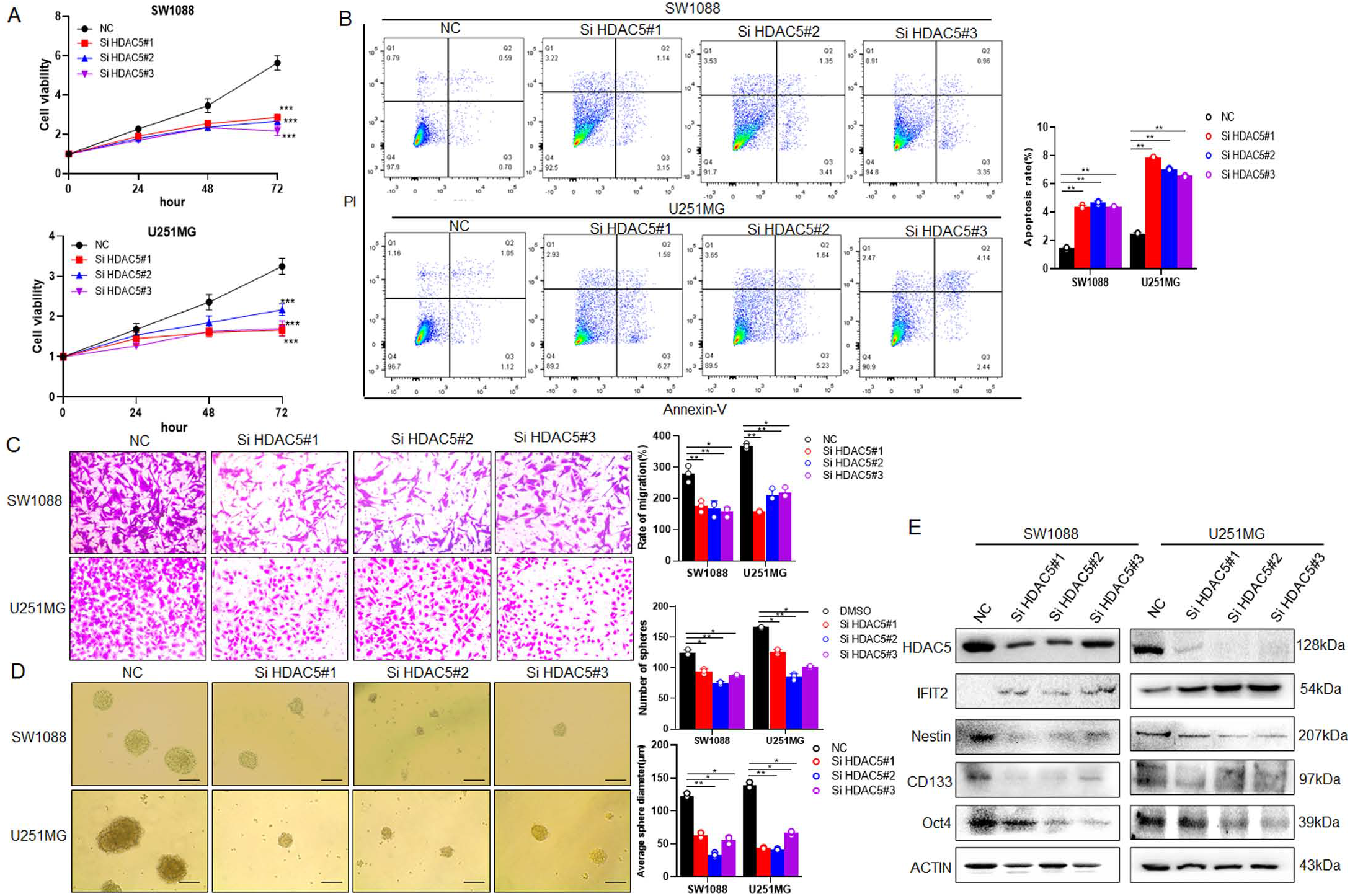
Overexpression of HDAC5 promotes the stemness and progression of glioma. SW1088 and U251MG cells were pre-transfected with SiRNA-NC or SiRNA-HDAC5 for 48h, **(A)** cell viability was determined using the CCK8 assay, **(B)** flow cytometric analysis was performed to investigate cell apoptosis, **(C)** the number of cells per field invading onto the lower surfaces of the filter was counted by invasion assay and **(D)** the number and the size of tumor spheres formed by SW1088 and U251MG cells was calculated. **(E)** The expression of HDAC5, IFIT2, Nestin, CD133 and Oct4 was detected by western blot after transfection with SiRNA-NC or SiRNA-HDAC5. Scale bar: 50 μm. Columns: mean of three replicates; *p<0.05, **p<0.01, and ***p<0.001.

### IFIT2 inhibits PKC pathway and suppresses tumor growth in vivo

It has been reported that IFIT2 involved in tumor inhibition mainly through decreasing the expression of TNF-α and phosphorylated AKT and PKC [16]. To investigate how IFIT2 suppress tumor progression of glioma, western blot was performed to analyze the potential pathway that IFIT2 regulated. As shown in Figure S8A, the expression of P-PKC was significantly reduced in IFIT2 overexpression cells. In addition, the expression of P-PKC was decreased in SiHDAC5 cells, which is consist with IFIT2 overexpression cells (Figure S8B). These data suggested that IFIT2 suppress glioma progression through inhibiting PKC pathway. Further, nude mice were inoculated subcutaneously with U251MG cells to conduct xenograft experiment. When tumors reached 60-80 mm^3^, LBH589 (20 mg/kg/day) or DMSO were administered intraperitoneally every 24 hrs for 5 days. The results showed that volume and weight of tumors was significantly inhibited by LBH589 treatment (Figure 7A-C). IHC demonstrated that LBH589 inhibited HDAC5 expression, and promoted the expression of IFIT2 (Figure 7D). Together, these data indicated HDAC5 could promote tumor growth by suppressing IFIT2 expression.

**Fig.7.**
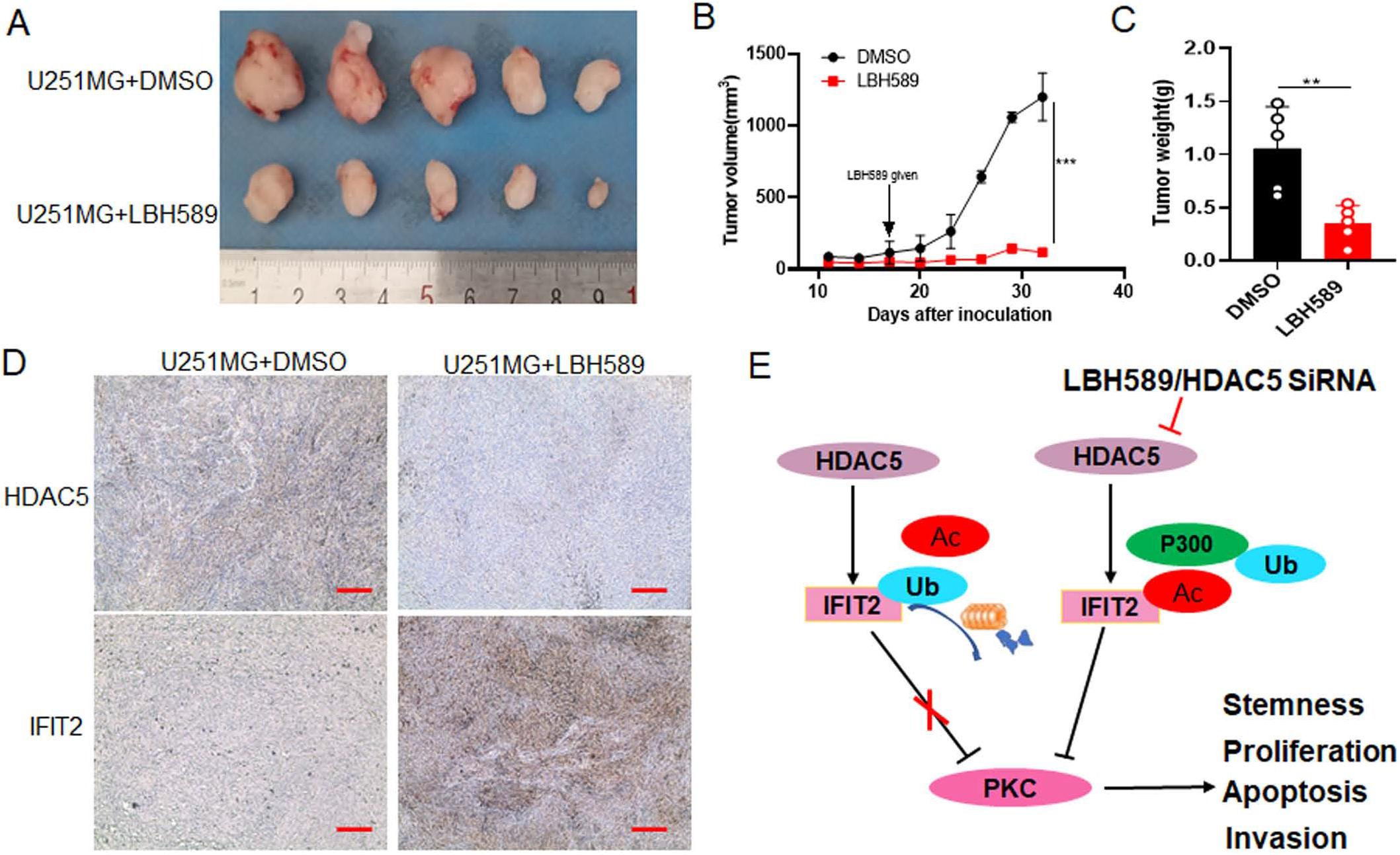
IFIT2 inhibits PKC pathway and suppresses tumor growth in vivo. **(A)** The glioma cell U251MG (1×10^7^) were injected into nude mice by subcutaneous injection. Mice were injected intraperitoneally with DMSO or LBH589 (20 mg/kg/day,5 times) when the tumor volumes were about 60-80 mm^3^. Representative images of the xenograft nodules are shown. **(B)** Growth curves of xenografts showed the mean volume of both treatment groups. **(C)** The weight of xenografts tumors was quantitatively analyzed. **(D)** The expression of HDAC5 and IFIT2 in the xenografts was analyzed by immunohistochemistry. Scale bar: 50 μm. **(E)** The model of the mechanism by which HDAC5 mediated IFIT2 deacetylation, increasing IFIT2 ubiquitination and inducing protein instability, thus promoting glioma stemness and progression is shown. **p < 0.01 and ***p<0.001.

## Discussion

Numerous studies indicated that IFIT2, a tumor suppressor gene, inhibited tumor growth, metastasis and induced cell apoptosis in many cancers like oral squamous carcinoma, colorectal cancer, leukemia, osteosarcoma and hepatocellular carcinoma [24]. In present study, we found that high expression of HDAC5, as well as low expression of P300 in glioma cells, decreased acetylation of IFIT2, then upregulated its ubiquitination and degradation, consequently promoting growth, metastasis and stemness of glioma. These result not only clarified a novel post-transcriptional regulatory mode of IFIT2, but also provide a new sight of molecular mechanism of HDACi in clinical application (Figure 7E).

Previous evidence has shown that IFIT2 was mainly transcriptionally regulated in cancer cells. Specifically, IFIT2, as an IFN-induced protein, was activated by IRF3 once the response element binding with regulator ISGF3 on the promoter of IFIT2, promoting cell apoptosis [25]. In gallbladder cancer, PLZF was found to significantly increase IFIT2 transcription by upregulating transcriptional factor STAT1 expression, which inhibited cell migration and invasion [26]. Downregulation of IFIT2 through Wnt/β-catenin pathway enables colorectal cancer cells resistant to apoptosis [27]. Moreover, STAT2 was identified to interact with P300 and CBP, activating the trans-domain and inducing histone acetylation, thus regulating the transcription of IFIT2 [19]. Notably, recent studies suggested that IFIT2 degraded in a proteasomal dependent manner, and blocking of proteasomal activity induced aggregation of IFIT2 in centrosomes, which promoted cell apoptosis [28]. Here, our findings showed that IFIT2 was not only suppressed at transcriptional level, but also directly downregulated by HDAC5 at post-translational level, which facilitated stemness and progression of glioma.

HDAC5, a member of class IIA HDACs, could translocated between the nucleus and cytoplasm during the dynamic and reversible acetylation modification, involving in many biological processes such as acting as an inhibitor of angiogenesis and regulating key angiogenesis-related gene expression in endothelial cells [29]. In addition, HDAC5 was also reported to mediate the deacetylation of SOX9, affecting its nuclear localization and participating in the progression of breast cancer [30]. Besides, HDAC5 could be recruited by TLX to the downstream target and inhibited its transcription, thereby regulating the proliferation of neural stem cells [31]. Our study demonstrated that HDAC5 mediated the deacetylation of IFIT2, while P300 was also involved in the acetylation modification of IFIT2, thereby inhibiting the expression of IFIT2 and promoting tumor progression of glioma.

A series of researches revealed that IFIT2 acts as a tumor suppressor gene. It promotes cell apoptosis through a mitochondrial pathway dependent on the role of Bcl2 protein [32]. IFIT2 descends angiogenesis and growth in OSCC cells through inhibiting TNF-α [15]. In gallbladder cancer, PLZF significantly promoted IFIT2 mRNA by increasing STAT1 protein level, and inhibited GBC cell migration and invasion through the EMT pathway [26]. Exhaustion of IFIT2 induced EMT, and further enhanced migration and invasion of OSCC cells mediating PKC signaling pathway [14]. Instead, overexpression of IFIT2 blocked oncogenic effects of BCR-ABL and downregulated Akt/mTOR pathway, thereby inhibiting the proliferation of CML cells and arresting the cell cycle in the G1 phase [16]. Our study uncovered that IFIT2 prohibited glioma stemness and progression by inhibiting PKC signaling.

In summary, we found that HDAC5 mediated IFIT2 deacetylation, increasing IFIT2 ubiquitination and inducing protein instability, thus promoting glioma stemness and progression. However, further investigation should be required such as the detailed modified site of IFIT2 in acetylation, and which E3 enzyme is involved in ubiquitination, and whether HDAC5 regulates the nuclear localization of IFIT2 remains unknown. Undoubtedly, our available data comprehensively identified a novel molecular mechanism that how IFIT2 was modulated in acetylation, and gave a sight in the role IFIT2 in glioma, which would be potentially beneficial to bench-to-bedside translation in glioma therapies.

## Authors’ contributions

L.Y. and D.F. performed study concept and design; Y.L., K.Z., X.P., Z.Z., and P.Z. performed development of methodology and writing, review and revision of the paper; Y.L., K.Z., and S.T. provided acquisition, analysis and interpretation of data, and statistical analysis; D.L. and L.S. provided technical and material support. All authors read and approved the final paper.

## Reference

1. Weller, M., et al., Glioma. Nat Rev Dis Primers, 2015. 1: p. 15017.

2. Bray, F., et al., Global cancer statistics 2018: GLOBOCAN estimates of incidence and mortality worldwide for 36 cancers in 185 countries. CA Cancer J Clin, 2018. 68(6): p. 394–424.

3. Wang, J., et al., BRD4 promotes glioma cell stemness via enhancing miR-142-5p-mediated activation of Wnt/beta-catenin signaling. Environ Toxicol, 2020. 35(3): p. 368–376.

4. Batlle, E. and H. Clevers, Cancer stem cells revisited. Nat Med, 2017. 23(10): p. 1124–1134.

5. Dawson, M.A. and T. Kouzarides, Cancer epigenetics: from mechanism to therapy. Cell, 2012. 150(1): p. 12–27.

6. Li, S., et al., Acetylation and Deacetylation of DNA Repair Proteins in Cancers. Front Oncol, 2020. 10: p. 573502.

7. Yoshino, J., et al., Loss of ARID1A induces a stemness gene ALDH1A1 expression with histone acetylation in the malignant subtype of cholangiocarcinoma. Carcinogenesis, 2020. 41(6): p. 734–742.

8. Bi, L., et al., HDAC11 regulates glycolysis through the LKB1/AMPK signaling pathway to maintain hepatocellular carcinoma stemness. Cancer Res, 2021.

9. McClure, J.J., X. Li, and C.J. Chou, Advances and Challenges of HDAC Inhibitors in Cancer Therapeutics. Adv Cancer Res, 2018. 138: p. 183–211.

10. Chen, R., et al., The application of histone deacetylases inhibitors in glioblastoma. J Exp Clin Cancer Res, 2020. 39(1): p. 138.

11. Pidugu, V.K., et al., Emerging Functions of Human IFIT Proteins in Cancer. Front Mol Biosci, 2019. 6: p. 148.

12. Nihira, N.T., et al., Acetylation-dependent regulation of MDM2 E3 ligase activity dictates its oncogenic function. Sci Signal, 2017. 10(466).

13. Wang, Y.P., et al., Regulation of G6PD acetylation by SIRT2 and KAT9 modulates NADPH homeostasis and cell survival during oxidative stress. EMBO J, 2014. 33(12): p. 1304–20.

14. Lai, K.C., et al., Depleting IFIT2 mediates atypical PKC signaling to enhance the migration and metastatic activity of oral squamous cell carcinoma cells. Oncogene, 2013. 32(32): p. 3686–97.

15. Lai, K.C., et al., Blocking TNF-alpha inhibits angiogenesis and growth of IFIT2-depleted metastatic oral squamous cell carcinoma cells. Cancer Lett, 2016. 370(2): p. 207–15.

16. Zhang, Z., et al., Overexpression of IFIT2 inhibits the proliferation of chronic myeloid leukemia cells by regulating the BCRABL/AKT/mTOR pathway. Int J Mol Med, 2020. 45(4): p. 1187–1194.

17. Koh, S.Y., et al., Baicalein Suppresses Stem Cell-Like Characteristics in Radio- and Chemoresistant MDA-MB-231 Human Breast Cancer Cells through Up-Regulation of IFIT2. Nutrients, 2019. 11(3).

18. Yao, Z.G., et al., LBH589 Inhibits Glioblastoma Growth and Angiogenesis Through Suppression of HIF-1alpha Expression. J Neuropathol Exp Neurol, 2017. 76(12): p. 1000–1007.

19. Paulson, M., et al., IFN-Stimulated transcription through a TBP-free acetyltransferase complex escapes viral shutoff. Nat Cell Biol, 2002. 4(2): p. 140–7.

20. Xue, Y., et al., HDAC5-mediated deacetylation and nuclear localisation of SOX9 is critical for tamoxifen resistance in breast cancer. Br J Cancer, 2019. 121(12): p. 1039–1049.

21. Taylor, P., et al., REST is a novel prognostic factor and therapeutic target for medulloblastoma. Mol Cancer Ther, 2012. 11(8): p. 1713–1723.

22. Narita, T., B.T. Weinert, and C. Choudhary, Functions and mechanisms of non-histone protein acetylation. Nat Rev Mol Cell Biol, 2019. 20(3): p. 156–174.

23. Ning, J.F., et al., Myc targeted CDK18 promotes ATR and homologous recombination to mediate PARP inhibitor resistance in glioblastoma. Nat Commun, 2019. 10(1): p. 2910.

24. Mamdani, H. and S.I. Jalal, Histone Deacetylase Inhibition in Non-small Cell Lung Cancer: Hype or Hope? Front Cell Dev Biol, 2020. 8: p. 582370.

25. Lee, J., et al., Absent in Melanoma 2 (AIM2) is an important mediator of interferon-dependent and -independent HLA-DRA and HLA-DRB gene expression in colorectal cancers. Oncogene, 2012. 31(10): p. 1242–53.

26. Shen, H., et al., PLZF inhibits proliferation and metastasis of gallbladder cancer by regulating IFIT2. Cell Death Dis, 2018. 9(2): p. 71.

27. Ohsugi, T., et al., Decreased expression of interferon-induced protein 2 (IFIT2) by Wnt/beta-catenin signaling confers anti-apoptotic properties to colorectal cancer cells. Oncotarget, 2017. 8(59): p. 100176–100186.

28. Lai, K.C., et al., IFN-induced protein with tetratricopeptide repeats 2 inhibits migration activity and increases survival of oral squamous cell carcinoma. Mol Cancer Res, 2008. 6(9): p. 1431–9.

29. Urbich, C., et al., HDAC5 is a repressor of angiogenesis and determines the angiogenic gene expression pattern of endothelial cells. Blood, 2009. 113(22): p. 5669–79.

30. Oltra, S.S., et al., HDAC5 Inhibitors as a Potential Treatment in Breast Cancer Affecting Very Young Women. Cancers (Basel), 2020. 12(2).

31. Sun, G., et al., Orphan nuclear receptor TLX recruits histone deacetylases to repress transcription and regulate neural stem cell proliferation. Proc Natl Acad Sci U S A, 2007. 104(39): p. 15282–7.

32. Reich, N.C., A death-promoting role for ISG54/IFIT2. J Interferon Cytokine Res, 2013. 33(4): p. 199–205.

