## Supplementary Figures for "Deacetylation of IFIT2 mediated by HDAC5 promotes the stemness and progression of glioma"

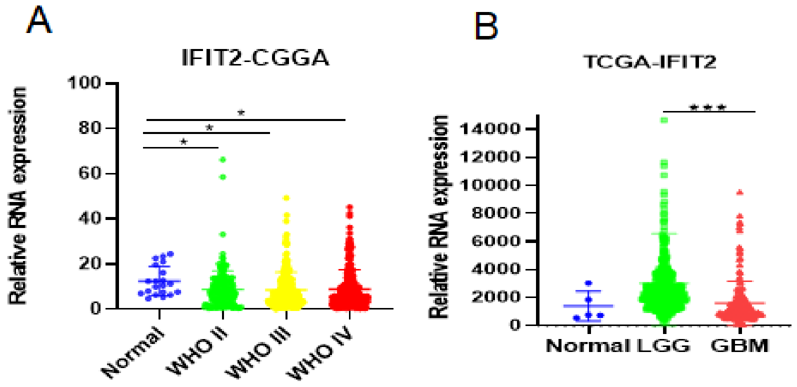


**FigS1.** The relative expression of IFIT2 in cancer tissues and their corresponding adjacent normal tissues based on data available from CGGA (A) and TCGA (B) database. *p<0.05, ***p<0.001. LGG: low-grade glioma, GBM: glioblastoma (high-grade glioma).

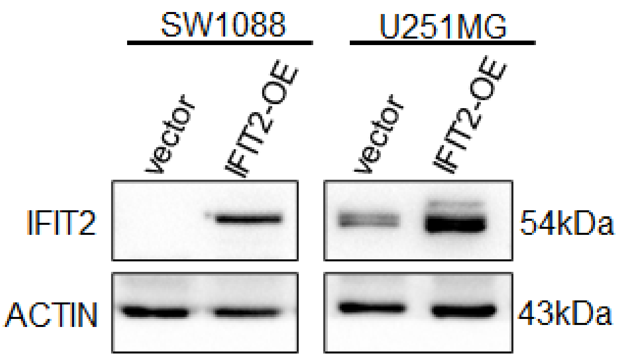


**FigS2.** SW1088 and U251MG cells were transfected with pcDNA3.1 3xFlag IFIT2 plasmid for 48h, and the expression of IFIT2 was analyzed by western blot.

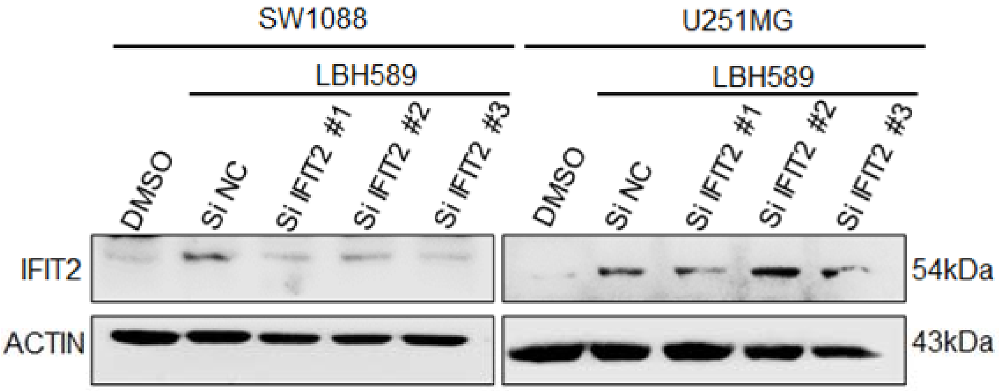


**FigS3.** After LBH589 (0.075μM) and transfection with siRNA-NC or siRNA-IFIT2 for 48h, and the expression of IFIT2 in SW1088 and U251MG was analyzed by western blot.

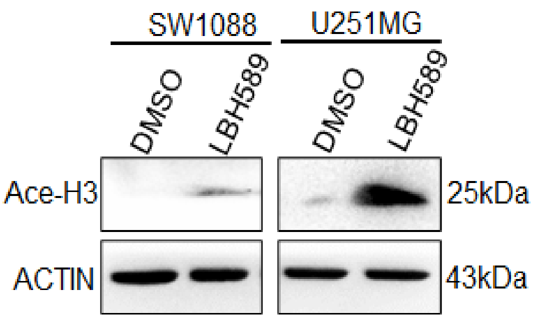


**FigS4.** The expression of ace-histone3 in SW1088 and U251MG cells with LBH589 treatment was analyzed by western blot.

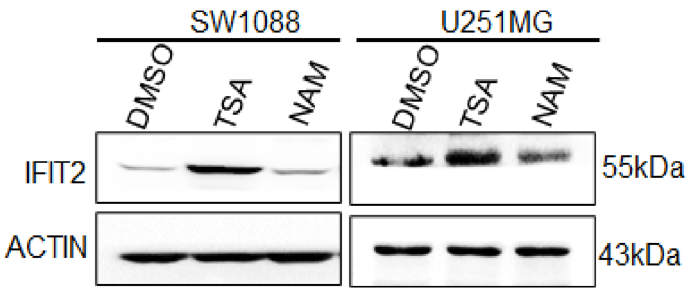


**FigS5.** Western blot analysis of IFIT2 protein in SW1088 and U251MG cells with pan-HDAC inhibitor (TSA, 1μM) or Sirtuin inhibitor (NAM,1mM) treatment for 48h.

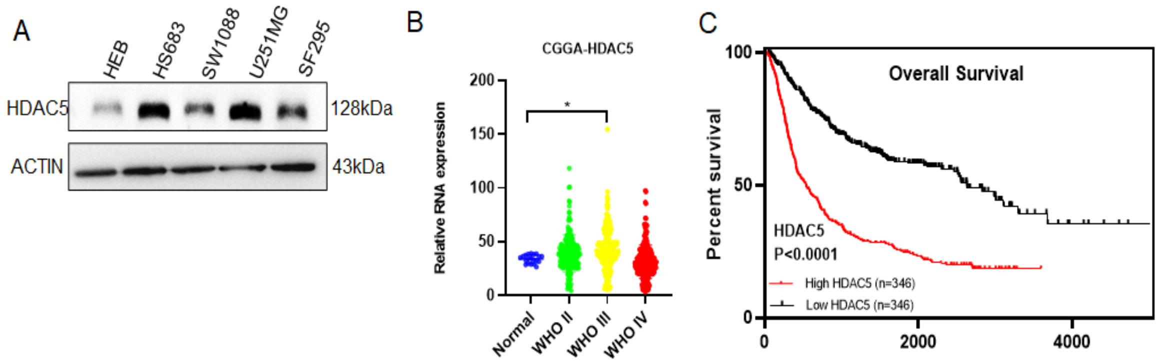


**FigS6. (A)** Protein expression of HDAC5 in glioma cells and normal glial cell HEB. **(B)** The relative expression of HDAC5 in glioma tissues and their corresponding adjacent normal tissues based on data available from CGGA database. *p<0.05. **(C)** Overall survival (OS) in patients with high (n = 346) vs. low (n = 346) levels of HDAC5 in glioma patients was plotted by the Kaplan–Meier method.

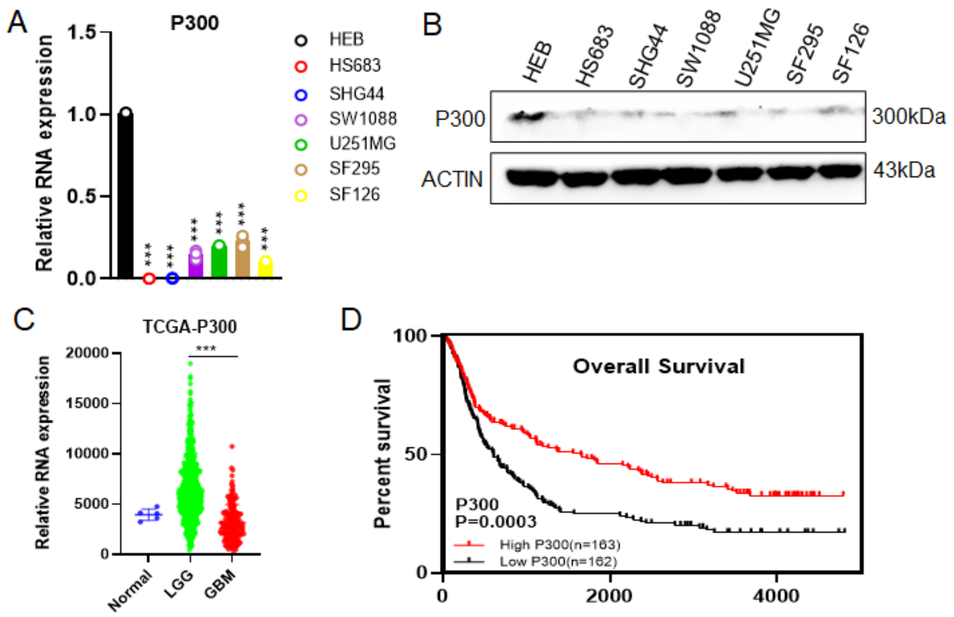


**FigS7. (A)** QPCR and **(B)**western blot were performed to detect P300 expression in glioma cells and normal glial cell HEB. ***p<0.001. **(C)** The relative expression of P300 in cancer tissues and their corresponding adjacent normal tissues based on data available from CGGA database. ***p<0.001. **(D)** Overall survival (OS) in patients with high (n = 346) vs. low (n = 346) levels of HDAC5 in glioma patients was plotted by the Kaplan–Meier method. LGG: low-grade glioma, GBM: glioblastoma (high-grade glioma).

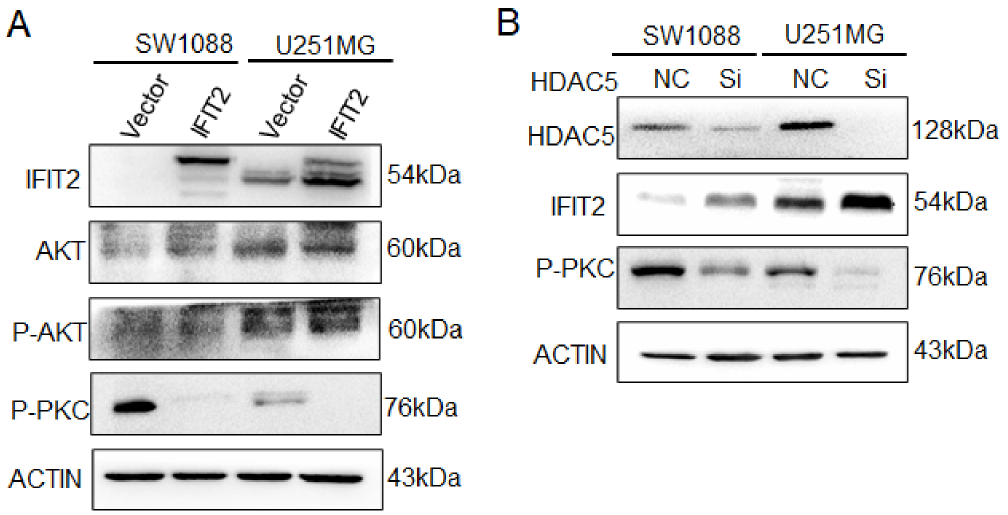


**FigS8. (A)** Western blot analysis for AKT, P-AKT and P-PKC in SW1088 and U251MG cells pre-transfected with IFIT2 plasmid for 48h. **(B)** Western blot analysis for IFIT2 and P-PKC in SW1088 and U251MG cells pre-transfected with siRNA-HDAC5 for 48h.
